# Deformation of membrane vesicles due to chiral surface proteins

**DOI:** 10.1101/2021.04.19.440377

**Authors:** Arabinda Behera, Gaurav Kumar, Sk Ashif Akram, Anirban Sain

## Abstract

Chiral, rod-like molecules can self-assemble into cylindrical membrane tubules and helical ribbons. They have been successfully modeled using the theory of chiral nematics. Models have also predicted the role of chiral lipids in forming nanometer-sized membrane buds in the cell. However, in most theoretical studies, the membrane shapes are considered fixed (cylinder, sphere, saddle, etc.), and their optimum radius of curvatures are found variationally by minimizing the energy of the composite system consisting of membrane and chiral nematics. Numerical simulations have only recently started to consider membrane deformation and chiral orientation simultaneously. Here we examine how deformable, closed membrane vesicles and chiral nematic rods mutually influence each other’s shape and orientation, respectively, using Monte-Carlo (MC) simulation on a closed triangulated surface. For this, we adopt a discrete form of chiral interaction between rods, originally proposed by Van der Meer et al. (1976) for off-lattice simulations. In our simulation, both conical and short cylindrical tubules emerge, depending on the strength of the chiral interaction and the intrinsic chirality of the molecules. We show that the Helfrich-Prost term, which couple nematic tilt with local membrane curvature in continuum models, can account for most of the observations in the simulation. At higher chirality, our theory also predicts chiral tweed phase on cones, with varying bandwidths.

PACS numbers: PACS : 87.16.-b, 87.15.Aa, 81.40.Jj, 87.15.Rn

## INTRODUCTION

Chiral molecules can self-assemble into membrane like structures. A broad class of chiral lipids, both synthetic [1, 2], as well as biological [3, 4], are known to self-assemble into membrane bilayers which take the shape of long cylindrical tubules and helical ribbons. The lipid orientations 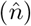 often exhibit constant tilt (*θ*) with respect to the local membrane normal 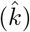 as in a smectic-C film. The projection of these tilted molecules (sin *θ*) on the cylindrical membrane surface shows helical striped pattern on tubules. These have been subject of many theoretical studies [5–9] which had employed theory of chiral nematics [10] to understand nematic textures on curved membrane surfaces. The so called *colloidal membrane* is another example where chiral, rod like virus particles, exhibiting strong depletion interaction among themselves, self assemble into a monolayers [11], but with a spatially nonuniform tilt. The monolayer can exhibit different shapes and curvatures depending on experimental conditions [11, 12].

While all these examples of self-assembled chiral membranes exhibit open structures with free edges, we will focus here on closed vesicles which are more constrained (by volume and area) in adjusting their shapes such that chiral interaction energy is minimized simultaneously. It has been proposed [13, 14] that such chiral lipid membranes could form spherical buds of very small size of order 50*nm*, although this has not been directly verified in experiments. However most of the studies on chiral membranes start with a predefined shape of the membrane (cylinder, ribbon, sphere, saddle etc) and look for the nematic texture that minimize the total energy of the system. Often the optimum curvature of the surface is computed variationally. For narrow, long tubes Ref[5, 9] had predicted uniform tubule radius *r* = 2*κ/c*_0_, where *κ* is the membrane bending modulus and *c*_0_ is the coupling strength between molecular chirality and the surface curvature. These initial theories had also predicted uniform tilt pattern of 45° with respect to the cylinder axis. In contrast, ref[7] predicted modulated tilt patterns. Recently ref[15] showed that such modulated patterns on uniform cylinders may not be constitute global minima of the system, instead modulated tilt patterns were shown to be stable on rippled cylinders. In contrast, here we use Monte-Carlo (MC) simulation to search for minimum energy configurations by varying both the vesicle shapes and the corresponding nematic texture simultaneously which opens up much wider possibilities in terms of shapes and patterns. Our analytic theory, in the second half of this paper, however, searches for optimum nematic texture on fixed shapes which are found in our simulations.

The scope of this study is even broader: the physics of uniformly tilted lipids is equally applicable to situations where an achiral membrane is decorated with rod-like chiral proteins which lie completely on the tangent surface of the membrane. Given that many membrane binding proteins are chiral in nature (e.g., proteins with BAR domains) their influence on the membrane shape is of interest. For example dynamin and ESCRT-III are well known filamentous proteins which form helical spirals and strongly modify membrane shapes. A recent study [16] have explored this angle; they used MD simulations to report shape transitions in cylindrical, open vesicles where chiral proteins bind to an achiral membrane.

Before describing the models let us note that chirality is essentially a three dimensional feature and therefore chiral order cannot be achieved if the rod-like molecules are lying on a flat surface. But if the surface is curved (deformed) then even if the rods lie on the local tangent plane of the curved surface it is possible to have a chiral order. Chirality implies that successive rods would like to maintain a preferred relative angle. This is however different from splay and bend. One good example is when the nematic rods are tangent to a helical space curve. This can be readily achieved when the nematics decorate the surface of a membrane tube.

### Model

The projection of the nematic rod, on the tangent plane of the membrane forms a 2D vector field **m**(**r**) of fixed magnitude *m* = sin *θ*. We will ignore this fixed magnitude and study the behavior of a unit vector field **m**(**r**). Following ref[6, 7, 13, 14, 17] the energy functional for chiral nematics is

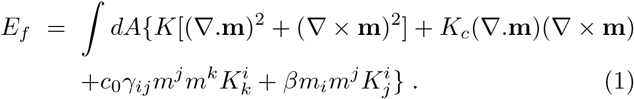

Here **m** is a 2D unit vector field in the tangent plane of the membrane. The first and the second terms are the splay and bend terms (with the same modulus *K*) [10]. The third term linear in ∇ × **m** promotes chiral order. In 2D, *Curl* **m** = *γ*_*ij*_*∂*^*i*^*m*^*j*^ is a pseudo scalar as 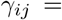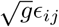, where *∈*_*ij*_ is the fully antisymmetric 2nd rank Levi civita tensor and *g* is the determinant of the metric tensor for the curved membrane surface. The invariant area differential *dA* also contains 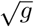 (see Eq-3 later). Further, *K*_*c*_ is the strength of this chiral interaction. Both the 4th and 5th terms couple the nematic field with the local surface curvature. In particular, the 4th term, known as the Helfrich-Prost term [5] is also chiral in nature. Next we will describe our discrete model which shares the symmetries of the above continuum model. Below we will also include the elastic energy terms for the pure membrane.

In the model below 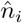 are the 3-D nematic directors located at the *i – th* vertex of the triangulated network and lies on the local tangent plane. The 3-D cross product 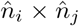 below is a pseudo-vector which contains the crucial chira effect.

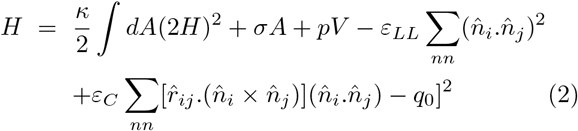

The first three terms are due to membrane bending (Helfrich), surface tension and pressure. The next two terms are discrete, involving only nearest neighbour (n-n) pairs Before describing the models let us note that chirality (*i, j*) on the triangulated network. 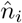 and 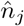 lie on the tangent planes of the n-n *i, j*-th vertices, and 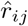 is the unit vector along 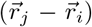, linking the *i, j*-th vertices. The *ε*_*LL*_ term (Lebwohl-Lasser interaction) accounts for splay and bend in the nematics and the last term, with modulus *ε*_*C*_, is the twist energy of the chiral nematics. The term linear in *q*_0_, emerging from the twist term, is the all important chiral term (a pseudo scalar due to the presence of a single cross product). It also couples local membrane curvature and the nematic tilt. For a flat membrane 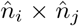 would be perpendicular to the plane while 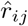 would lie on the plane, resulting in zero contribution from the dot product. Note that the nematic interactions in the continuum Eq.1 had two chiral terms (with coefficients *c*_0_ and *K*_*c*_) where as here, in Eq.2, we have only one with coefficient *ε*_*C*_ *q*_0_.

In principle, the dot and the cross products between the neighboring vectors 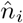 and 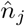, in Eq.2, should be carried out after parallel transporting 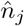 to the tangent plane of 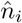 or the converse. But note that for a given vertex *i* there are typically five to six nearest neighbors *j* and all the 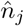 s’ can be parallel transported to the tangent plane of 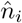. Similarly for *n*_*j*_ all its neighbors can be transported to *n*_*j*_’s tangent plane. In this symmetric scheme all (*i − j*) pairs would be counted twice. However in practice this scheme fails for the cross product terms. This is because for a reasonably triangulated smooth surface, the tangent planes of the neighboring vertices are almost parallel to each other. Further, 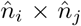 are very small (since the nematic field varies smoothly across neighboring vertices) and their vector directions (away from the vesicle or into the vesicle) are also very sensitive to the discretization errors. Note that the tangent plane at a vertex is calculated by averaging over the five or six neighboring triangles [18]. Furthermore, the neighboring tangent planes being close, 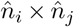 is almost perpendicular to 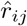, which is the unit vector connecting the vertices *i* and *j*. This makes the contribution of 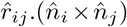 even smaller and fluctuating in sign. While summing over five to six such small numbers with random signs, often the noise to signal ratio is too high to capture any systematic chiral effect. In contrast, when the simulation was run without parallel transport, significant chiral effects could be captured. The calculation of 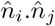 terms, being close to one and positive, do not suffer from this problem and gives nearly the same result with and without parallel transport.

Therefore we carried out all the dot and the cross products without parallel transport. We note that when the discretization is reasonably good the local tangent planes of vertices *i* and *j* are close to parallel. This makes the effect of parallel transport minimal, but more importantly absence of parallel transport reduce the numerical noise involved in the parallel transport operation over nearly parallel planes. To verify the validity of this scheme we reproduced all the standard chiral and achiral nematic patterns known for spherical and cylindrical surfaces. For example, a) for rigid sphere, with *ε*_*C*_ = 0 in eq.2, we reproduced the standard equilibrium configuration of nematics on a rigid sphere, i.e., four +1/2 defects located at the vertices of a tetrahedron, b) for a rigid sphere, by imposing two +1 defects at the north and south poles we reproduced the standard ‘S’ shape configuration [14], for nonzero *ε*_*C*_, with *ϕ* ≃ 45° (see Fig-2), and c) for cylinder, see Fig3-b,c,e and f) for various boundary conditions at the two ends, when *q*_0_ is not too small, we recovered the standard *ϕ* ≃ 45° configuration [14] in the bulk. For very small *q*_0_ values (see Fig3a and d), we obtained uniform orientation along the axial or the azimuthal directions, depending on the boundary conditions.

The simulation results presented here are for triangulated network forming closed surfaces consisting of *N* vertices. For rigid spherical surfaces we used *N* = 677 vertices and for the deformed vesicle we used *N* = 2030 vertices. *ε*_*LL*_ and *ε*_*C*_ have dimensions of energy and we present results for dimensionless parameters *ε*_*C*_/*ε*_*LL*_ and *q*_0_. As mentioned earlier the nematic term is borrowed from the standard Lebwohl-Lasher model [19] for nematics with nearest neighbour interactions. This term accounts for both splay and bend distortions within one Frank constant approximation. It has the local inversion symmetry 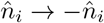 required by uniaxial nematics.

## RESULTS

Before presenting our main results on the vesicle deformations due to chiral nematics and the patterns of nematic arrangements on them, we first verify various ground state nematic textures, on spherical and cylindrical surfaces, that are well known in the literature [9, 13, 14].

### Chiral nematics on rigid surfaces

#### Sphere

Equilibrium nematic texture on a rigid vesicle is well known. Four +1/2 defects form and they try to maintain equal distance and as a result the defects arrange on a symmetric tetrahedron. When the nematic rods are chiral we find that only at low *q*_0_ value the nonchiral defect arrangement (i.e., four +1/2s’) are recovered (see Fig.1). As *q*_0_ is increased population of both +1/2 and −1/2 defects increase. This is due to the fact that increase in *q*_0_ increases the relative tilts between the neighbouring directors and as a result director arrange themselves on nearly closed loops. Radius of these loops decrease with increasing *q*_0_. Also note that the reflection symmetry of the texture around a +1/2 defect is broken when *q*_0_ is non-zero.

**Figure 1:**
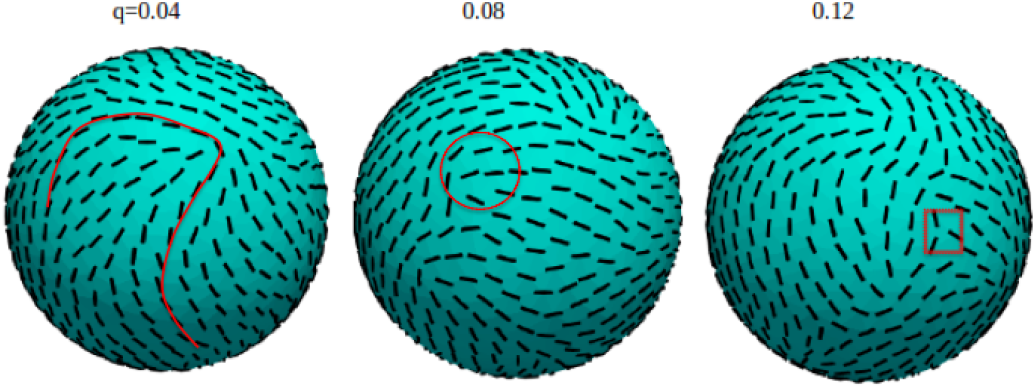
As intrinsic chirality *q*_0_ increases (left to right) number of defects increases. At low *q*_0_ (left most figure) only four +1/2 defects can be seen which is the lowest energy defect structure for a sphere. As defects repel each other elastic energy due to nematic distortion increases with population of defects, but at higher *q*_0_ this cost could be offset by the energy gain from the chiral term. We have indicated the core of a +1/2 and a −1/2 defect by a red circle and a square, respectively. On the left most figure we mark a loop of nematics around a +1/2 defect. Note that the loop does not have reflection symmetry. At higher chirality as more +1/2 defects form −1/2 defects also appear in order to to bring down the total charge to +2 (Poincaré’s index theorem). Parameter : *ε*_*c*_/*ε*_*LL*_ = 200.

In Ref[14] a plausible nematic texture was proposed for the bud structure (see Fig.2) with two centres of chirality at the top and the bottom. They also performed variational calculation to derive the energy of such a texture. Having imposed two +1 defects at the two poles, using our MC simulation on rigid sphere, we could verify this as the lowest energy configuration at sufficiently small *q*_0_.

**Figure 2:**
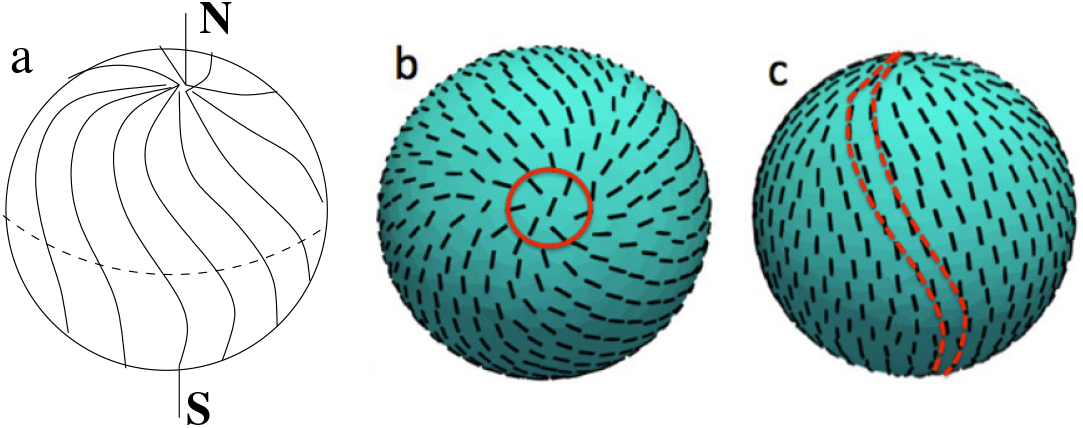
Chiral S-like patterns with two +1 defects imposed at the two poles of a spherical vesicle. a) Shows the low energy nematic texture for a bud which is connected to the base of a membrane vesicle by a narrow neck as was proposed in Ref[14]. This schematic was taken from their paper. (b) and (c) are the results from our simulation. (b) shows the top view and (c) the side view of the nematic texture. Orientations of the nematics inside the red circle were held fixed to impose the +1 defects (and similarly at the opposite pole). In the side view (c), approximate S-like patterns can be seen (see two successive red dashed lines). A perfect S-like pattern requires that number of latitude lines originating from the poles have to be same as the ones crossing the equator. This requires that the density of nematics have to be high near the poles. This is not the case in our triangulation scheme. We have roughly uniform density of vertices on the sphere. As a result we have more number of latitudes crossing the equator than that coming out from the pole. Parameters : *q*_0_ = 0.008 and *ε*_*c*_/*ε*_*LL*_ = 200.

#### Cylinder

The nematic arrangements on a cylindrical surface has been studied theoretically by various authors[8, 14, 15]. In ref.[14], the authors used symmetry arguments to arrive at the conclusion that the nematic rods must form a uniform tilt pattern on a cylinder, to break the reflection symmetry. They argued, in order to break the reflection symmetry, the nematic rods must form an angle between 0° and 90° with respect to the equator. So, the simplest of such configuration will be a pattern with uniform angle all over the cylinder. In this section, we discuss the uniform tilt pattern on a rigid cylinder using numerical simulations(see Fig.3).

**Figure 3:**
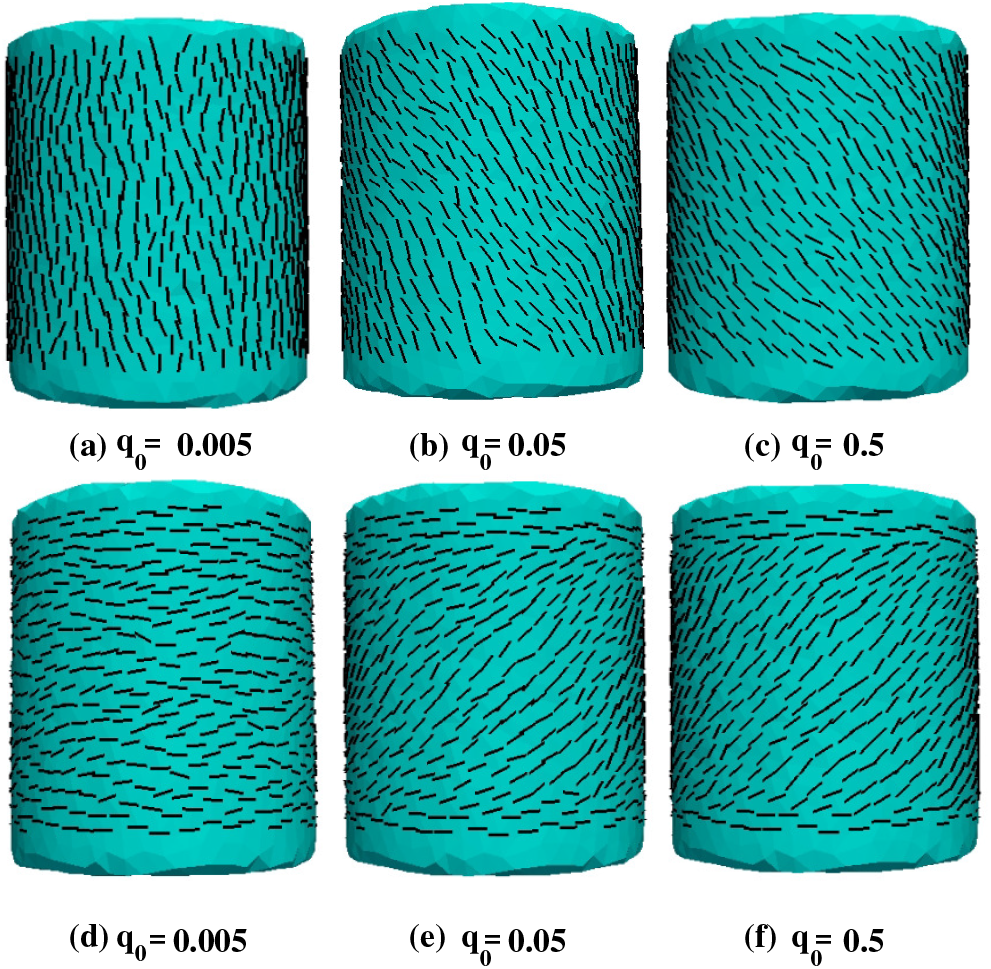
Chiral nematic patterns on a fixed cylinder with increasing intrinsic chirality (*q*_0_). In (a)-(c), no boundary conditions were imposed on the nematic orientations at the top and bottom. In (d)-(e), the nematic directors were held fixed along the azimuthal direction at the top and bottom. For both cases, 45° minimal orientation were attained in the bulk when chirality was not too small. At very small chirality uniform axial or azimuthal orientation was obtained.

For our simulations, we considered a triangulated cylinder with nematic rods distributed uniformly from the base to the top of the cylinder. We performed the simulations with a fixed cylinder for two cases: i) without any boundary condition, ii) with fixed boundary conditions imposed at the top and bottom two layers of the nematic arrangements. For the first case, at low *q*_0_, we found the nematic almost aligned parallel to each other making an angle close to 90° with respect to the equator(see Fig.3a). This configuration does not break the reflection symmetry as it does not have a distinct mirror image, hence it is a non-chiral arrangement. However, as we increased the *q*_0_’s value, we observed a uniform tilt phase, where the nematic rods make an uniform angle(0° < Φ < 90°) with respect to the equator(Fig.3(b-c)). However, a further increase in *q*_0_’s value beyond *q*_0_ = 0.5, forms similar configuration of uniform tilt pattern, which are not qualitatively distinct from each other. However, these patterns satisfy a common criteria- they break the reflection symmetry.

In the second case, we imposed a fixed boundary condition at the two top and bottom layers of the cylinder. The nematic rods at these layers were kept fixed in simulation, however, the bulk nematic rods were allowed to vary. At these boundary layers, the nematic rods were aligned along the azimuthal-direction of the cylinder, i.e., they make an angle 0° with the equator. For low *q*_0_, we observed the nematic rods almost lie along the equatorial-direction, in accordance with the boundary conditions imposed at the boundaries of the cylinder (Fig.3d). However, as we increase the *q*_0_’s value, we recover the uniform tilt pattern in the bulk part of the cylinder(Fig.3d-e). Near the two boundary layers at top and bottom of the cylinder, the nematic rods try to mimic the boundary condition. However, as we move towards the bulk part, the nematic rods slowly deviate from the imposed boundary condition and try to form a uniform tilt pattern, breaking the reflection symmetry. Similar to the previous case, the tilt patterns for higher *q*_0_ does not produce qualitatively distinct patterns. So, our simulations show the effects of chirality on the system in a narrow range of *q*_0_ value. We did not observe modulated tilt patterns as proposed in ref.[7].

### Chiral nematics on deformable vesicles

Chiral nematic rods can attain their intrinsic chirality easily on curved surfaces in order to minimize the contribution of the chiral term(s) in the hamiltonian. As a result we saw emergence of chiral rolls of increasingly smaller radii, on rigid spherical surfaces, as *q*_0_ was increased. This occurred even at the cost of proliferation of defects (see Fig.1). When the vesicle is soft and deformable it is interesting to ask what type of deformations will emerge which will minimize the full hamiltonian (Eq.2) consisting of the membrane energy (the first three terms) and the nematic energy costs (the rest of the terms). In Fig.4 we show the result of our simulations for chiral nematics on deformable vesicle. What cathces the eyes easily are the conical and cylindrical tubules at different values of *q*_0_ and *∈*_*c*_/*∈*_*LL*_, and the resulting chiral arrangements of the nematics around these tubules. The sub-figures in the 3 × 3 panel, can be broadly classified in three categories according to the value of the product *p* = *q*_0_.*∈*_*c*_/*∈*_*LL*_, which is essentially the strength of the chiral term in the discrete hamiltonian in Eq.2. a) When *p* is moderate, i.e., for sub-figures a, e and i, on the diagonal, the tubule lengths are moderate. b) When *p* is relatively smaller i.e., for the lower left triangle (consisting of d, g, and h) the tubules are wider and longer. c) When *p* is relatively high, i.e., for the upper right triangle (consisting of b, c, and f) the protrusions are relatively narrower and smaller. At these high values of *p* we also reach the limit of our resolution and and smooth spirals cannot be identified as the relative angles between the neighboring nematics are high and the arrangements look somewhat random. Note that for small *p* the nematics exhibit large helical pitch on the tubules and point nearly towards the tip of the tubule (see Fig.5a and Fig.6a), where as for moderate *p* the helical pitch in the spirals are smaller (see Fig.6b) and the nematics can be seen to be spiraling around the tubule (see Fig.5b,c). Note that for extremely high values of *p*, see Fig.4c, the protrusions appear to be small incomplete membrane buds, although the nematic arrangement on them is unclear. Interestingly, Ref[14], using continuum theory, had predicted formation of small membrane buds due to chiral effects. In contrast, at relatively low *p* wide, tube like membrane undulations form, see Fig.4-g, but instead of protruding out like tubules they cause large membrane invaginations or concave valleys. We do not have any quantitative understanding of these structures and their instabilities yet.

**Figure 4:**
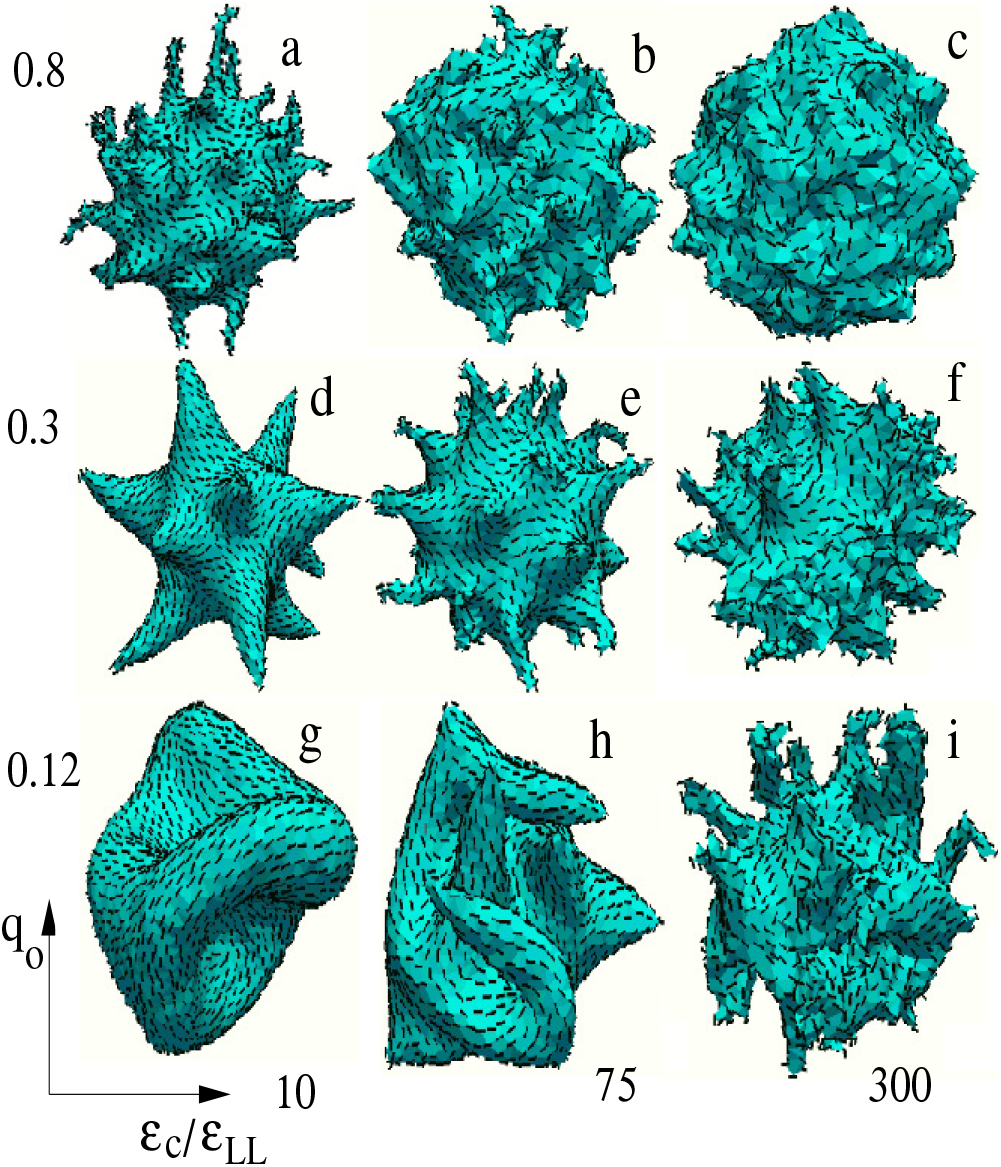
Shape deformation of a vesicle due to chiral nematics. Shapes are shown as a function of the scaled twist modulus *∈*_*c*_/*∈*_*LL*_ and intrinsic chirality *q*_0_. Simulations are done at constant pressure (*P*) and surface tension (*σ*). Therefore volume and area are not conserved across these different shapes. Conical and cylindrical protrusions of varying radius and lengths are seen at different parameter values. As intrinsic chirality (*q*_0_) is reduced (down the columns) chiral rolls with larger pitch form on wider cones as the inter-nematic inclination angle change over large length scales. When both *q*_0_ and *∈*_*c*_/*∈*_*LL*_ are high, rounded protrusions form, which change to cylindrical tubules at smaller *q*_0_. Cylindrical protrusions dominate at low *q*_0_. When *∈*_*c*_/*∈*_*LL*_ is reduced at low *q*_0_, wide cylindrical deformations occur. Other parameters: *κ* = 10, *σ* = 0.5.

**Figure 5:**
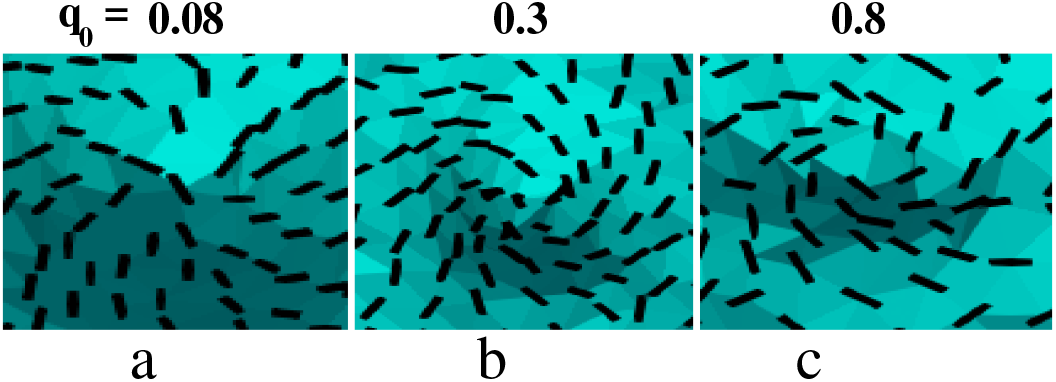
A Closer look at the chiral rolls around partial buds, tubules and wide membrane invaginations. These shapes are from column-2 of Fig.4. At low intrinsic chirality, no chiral rolls observed (*q*_0_ = 0.08). However, with increase in intrinsic chirality *q*_0_, the rolls became more prominent with larger angular variation at each successive layer.

**Figure 6:**
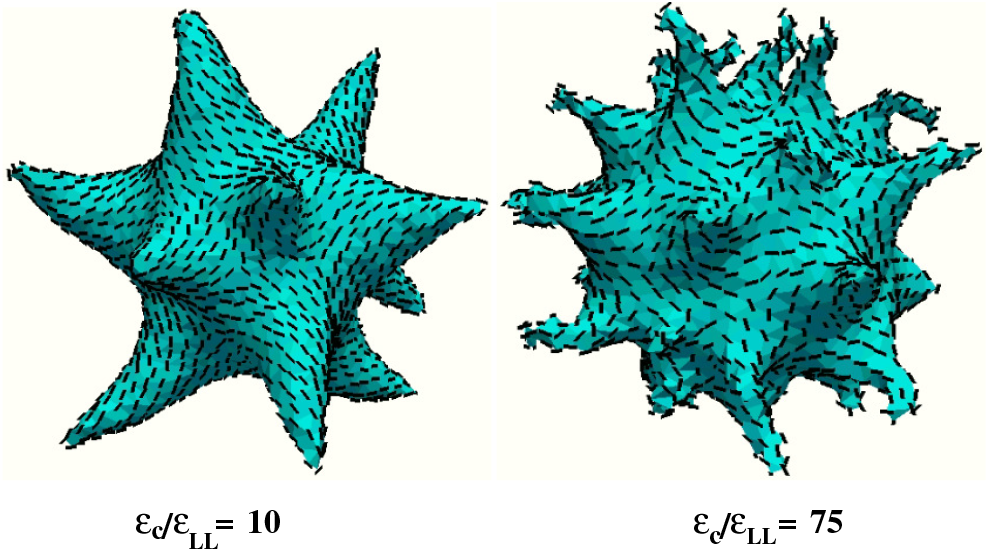
Chiral nematic patterns on deformed vesicles obtained from simulation for same intrinsic chirality *q*_0_ = 0.3 but different *ε*_*c*_/*ε*_*LL*_ ratio. In both cases, we observed cone-like protrusions.

However it is not possible to classify all these structures using only one parameter *q*_0_*∈*_*c*_/*∈*_*LL*_. The hamiltonian also contains a term proportional to *∈*_*c*_/*∈*_*LL*_, which is non-chiral but still contributes to the total energy. Motivated by the formation of distinct conical and cylindrical tubules from closed vesicles, we examine the possible arrangements of the chiral nematics on these shapes, using continuum theory. These tubules typically exhibits an aster type +1 topological defect at the tip and as a result nematic directors splay away from the tip. We will use this feature as a boundary condition.

## THEORY : NEMATIC ORDERING ON CONE AND CYLINDER

We now use the continuum Frank free-energy, given in Eq.1 (reproduced below) to study the arrangement of chiral nematics on conical and cylindrical tubules [14]. Here we have omitted the non-chiral curvature coupling term (with coefficient *β*),

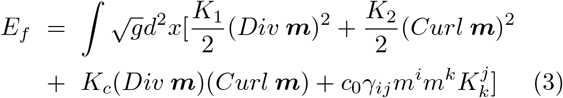

We define the local tangent plane on a conical surface (of height *H* and semi angle *α*), by unit vectors 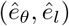. Here 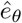 is the standard azimuthal vector in cylindrical coordinate system and 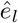 is orthogonal to 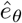 and points away from the apex. The surface is represented in the cylindrical coordinate system using the position vector 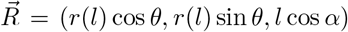. Here *l* is the distance from the apex of the cone and *α* is the semi-angle. From Fig.7, *r*(*l*) = *l* sin *α*, and *z* = *l* cos *α*. The nematic lies in the tangent plane making an angle Φ with 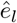. Φ depends on only *l* as we assume azimuthal symmetry.

**Figure 7:**
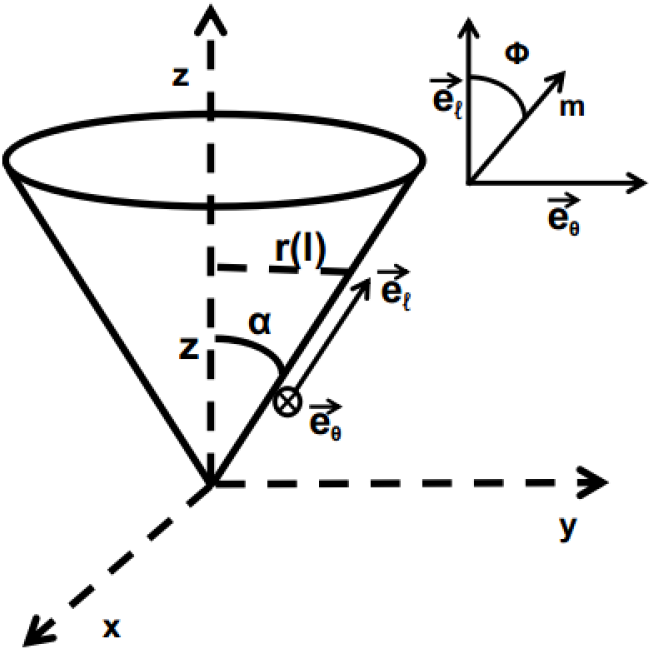
Orthogonal basis vectors 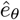 and 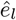 on the tangent plane of a cone, with semi-angle *α* and height *H*. Inset shows the nematic director 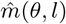 making an angle Φ with 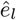 in the tangent plane.

Here, *Div **m*** and *Curl **m*** represent the covariant divergence and curl on the conical surface, and *g* is the determinant of the corresponding metric tensor *g*_*ij*_ of the curved surface. All these are computed explicitly in Appendix, for a conical surface. The first and the second terms here represent the generalized splay and bending, respectively. The third and the last terms are chiral in nature, i.e., they change sign under reflection (Φ ↦ −Φ). The antisymmetric tensor 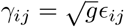, ensures the chirality of the last term, and its nonzero off-diagonal elements are 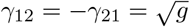 and *γ*_11_ = *γ*_22_ = 0.

The Euler-Lagrange equation for Eq.3, for a cone, is given in the Appendix (see Eq.14). We solve the non-linear differential equation (Eq.14) numerically, using Mathematica. To mimic the nematic directors splaying away from the apex we impose the boundary conditions Φ(*l* = 0) = 0, i.e., directors pointing towards the apex. To avoid the singularity at the apex we impose this boundary condition infinitesimally away from the apex, i.e., Φ(*l* = *δ* → 0). For the closed vesicles which we simulated, the base of the conical protrusions are attached to the main body of the vesicle where the nematic orientations are not precisely known. Typically, this is the starting point of a spiral of unknown pitch. We therefore impose torque free boundary condition [20] at the base of the cone, at *l* = *H/* cos *α*, so that the pitch is not constrained by this boundary condition. It amounts to 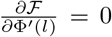, where Φ′ implies derivative with respect to *l* and 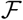 is the energy density. We discuss the solutions next.

We first study the effects of the chiral Helfrich-Prost term. We fix the value of *K*_*c*_ = 0.1, and vary only *c*_0_. The results are shown in Fig.8. At small *c*_0_, the director angle Φ, increases monotonically from 0°, at the vertex, to close to *π*/2 at the base of the cone. However, for a larger *c*_0_ values, Φ oscillates with *l* with an increasing wave length as the radius *r*(*l*) increases. This gives rise to band like patterns similar to tweed texture [14, 21], but with variable band width. Band-like modulated tilt patterns (with uniform band width) have been predicted theoretically on cylindrical surfaces by Selinger et al.[9]. The larger variation of Φ at higher *c*_0_ is forced by the stronger chiral interaction requiring relatively larger angle between the neighboring directors. This is equivalent to shorter pitch in the spirals, obtained in our simulations, at larger *q*_0_*∈*_*c*_/*∈*_*LL*_ values. As a result, the nematic directors rapidly change their orientation in subsequent layers (away from the apex) before reaching the boundary, producing multi-valued solutions of Φ. However our simulation does not produce multiple bands on the conical tubules; there only the pitch increases with *q*_0_*∈*_*c*_/*∈*_*LL*_, similar to the low *c*_0_ structures produced here in the continuum theory. We also found, that at low *c*_0_ values, variation of *K*_*c*_ does not have much effect (figure not shown). However at higher *c*_0_, increase in *K*_*c*_ can give rise to banded tweed-like nematic texture of increasing width, see Fig.9.

**Figure 8:**
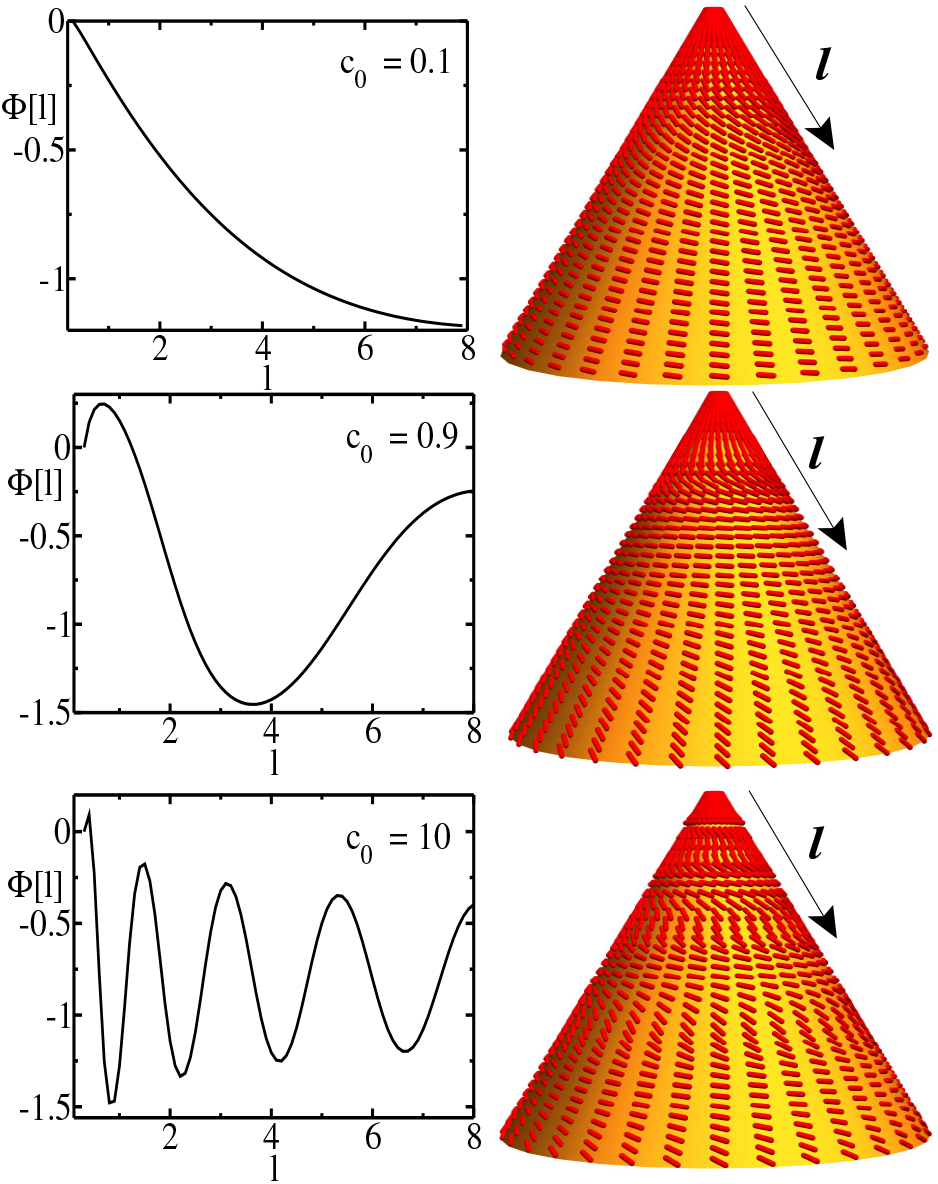
Left panel: Variation of Φ(*l*) with *l* for fixed *K*_*c*_ and changing *c*_0_. Right panel: Nematic (red) on a cone (yellow surface). The Helfrich-Prost constant *c*_0_ is in increasing order from top to bottom (see legends). Higher *c*_0_ values induce modulated tilt-patterns on the cone’s surface.

**Figure 9:**
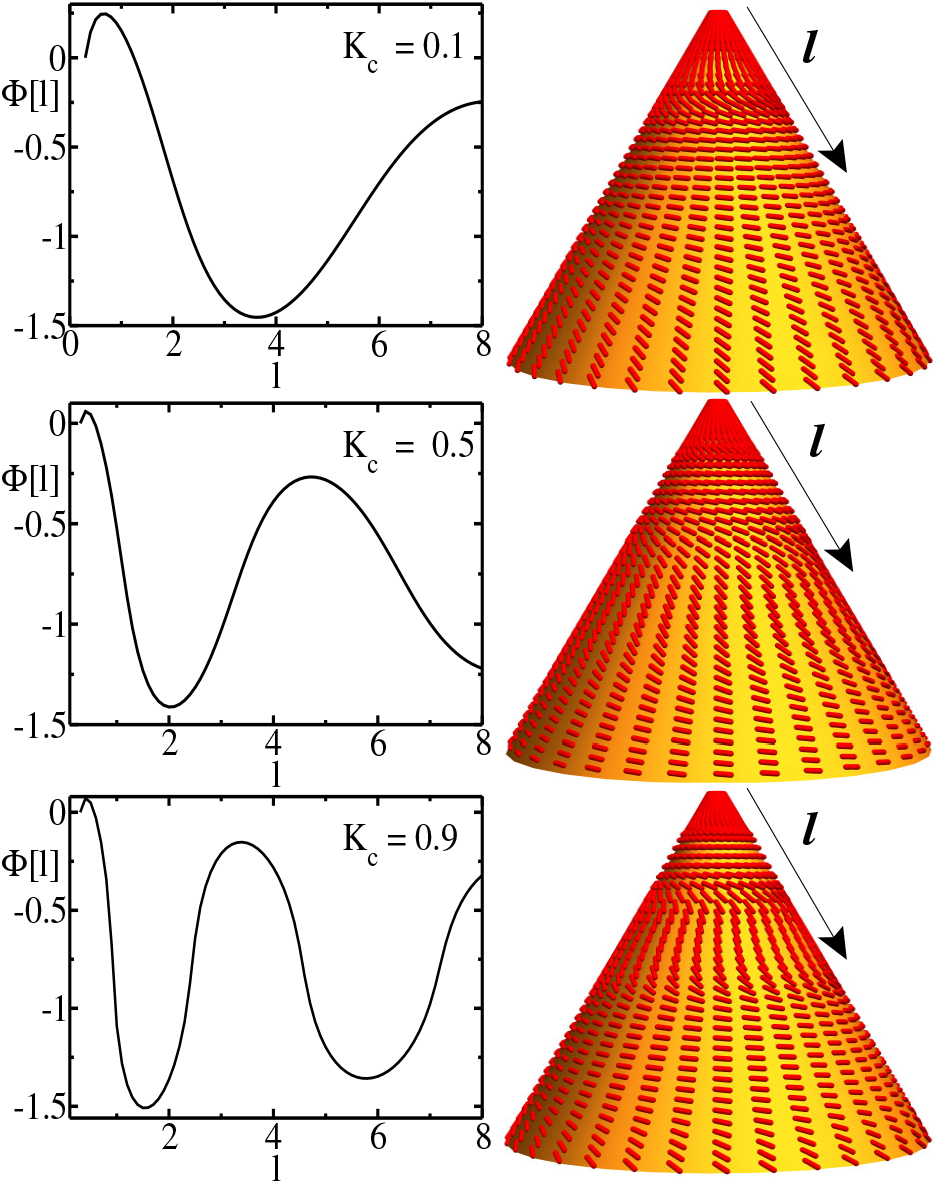
Left panel: Variation of Φ(*l*) with *l* for fixed *c*_0_ = 0.9 and changing *K*_*c*_. Right panel: shows corresponding nematic arrangements (red line segments) on a cone (yellow surface). *K*_*c*_ increases from top to bottom (see legends). Higher *K*_*c*_ values induce modulated tilt-patterns on the cone’s surface.

Next we look for chiral nematic patterns on a cylindrical surface of height *H* and radius *r*. The Euler-Lagrange equation for a cylinder is derived in the appendix (see Eq.18). To solve Eq.18, we imposed the same boundary conditions as in the case of a cone, i.e., Φ(0) = 0 and torque free boundary at *H*. The results for the cylinder are shown in Fig.10. Here, we kept *K*_*c*_ fixed and varied only the *c*_0_ value. Monotonic increase of tilt angle Φ is seen at small *c*_0_ values, with higher pitch at relatively higher *c*_0_ value. Uniform tweed-texture is obtained at high *c*_0_ value, see Fig.10. Here Φ(*z*) oscillates in the range [0, *π*/2] in a regular fashion.

**Figure 10:**
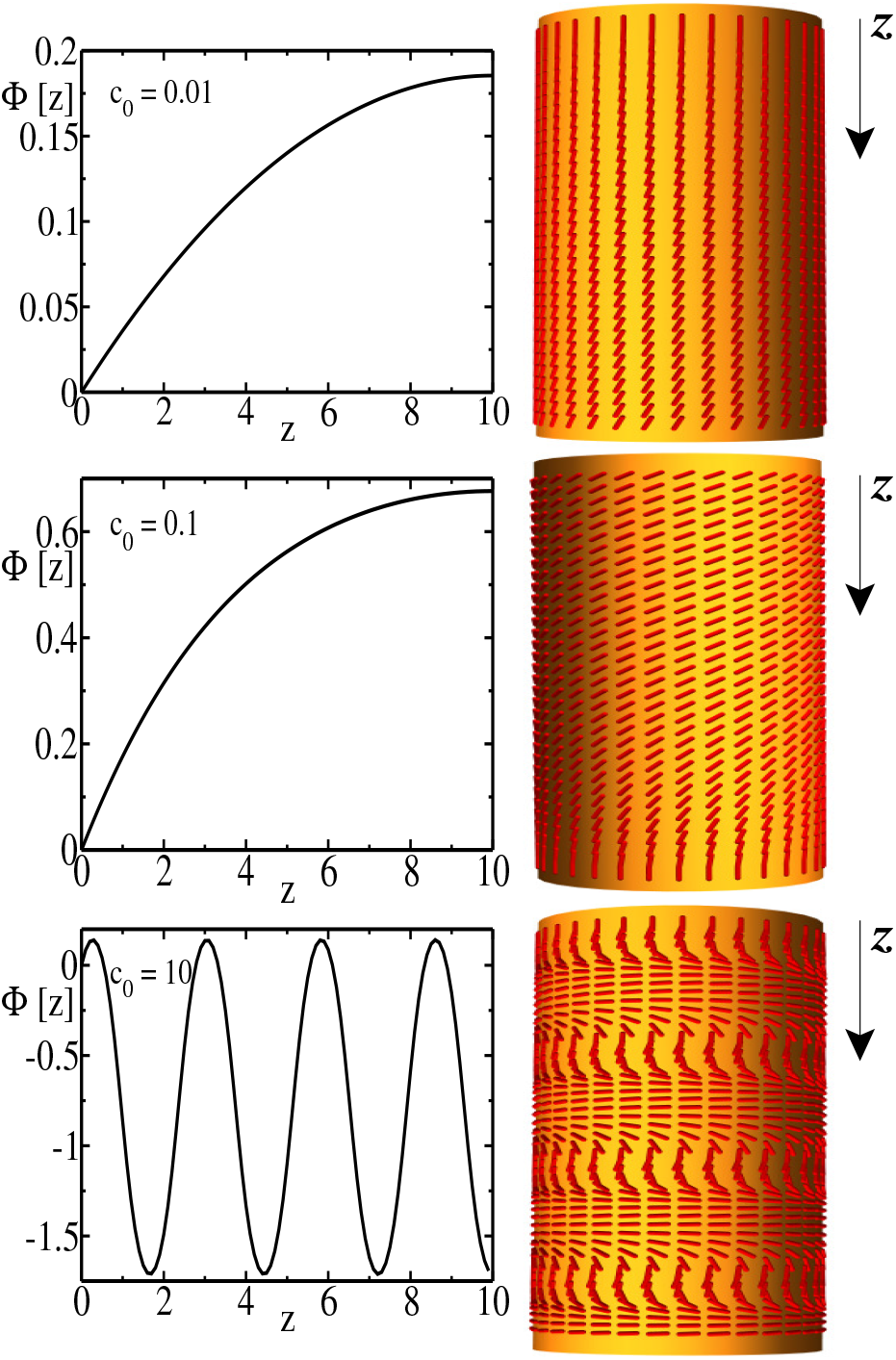
Left panel: Variation of Φ(*z*) with *z*. The plots are in increasing order of *c*_0_ from top to bottom. Right panel: the corresponding nematic arrangement(red tube) on cylinder’s surface(yellow). Here, *K*_*c*_ = 0.1, height of cone *H* = 10. A higher *c*_0_ produces modulated tilt patterns on the cylinder, which are qualitatively similar to the tweed texture discussed in ref.[14].

## DISCUSSION

We will now discuss what is new that we learn from this work. It is well known that chiral order is frustrated in the bulk of a flat surface but can be easily satisfied on curved surfaces or at surface boundaries. This is because the nematic rods can access greater angular space to maintain relative tilt with their neighbors. This effect is manifest in certain chiral lipids that self assemble into long cylindrical tubules. However, it was not obvious that even closed vesicles can overcome bending energy costs and grow protrusions (conical or cylindrical) due to chirality. The size of the protrusion depend on twist modulus and intrinsic chirality. Using a discrete model of chiral nematics on a triangulated closed surface we demonstrated this effect. We validated our model and the numerical scheme by verifying the standard chiral patterns on rigid spheres and cylinders, predicted in the literature by continuum models. While the standard continuum model that we considered had two chiral terms (the Helfrich-Prost term and the *K*_*c*_ dependent term which is allowed by the inside-outside assymmetry of a closed vesicle), our discrete model had only one chiral term. In the continuum theory we had ignored non-chiral curvature coupling terms 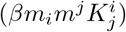, anisotropic curvature terms and other terms higher-order in ***m*** allowed by symmetry [14, 22]. Our texture calculation showed that between the two chiral terms the Helfrich-Prost term turned out to be more dominant in promoting the modulated tilt patterns on surfaces of cones and cylinders. However our modulated tilt pattern on cylinder is probably a metastable structure [15] which is stabilized due to our boundary condition Φ = 0, at *z* = 0.

Within the resolution of our simulation we did not observe clear bud-like membrane protrusion as predicted in ref[14] for chiral nematics. There could be two reasons for this: a) in ref[14] the emergence of spherical bud required positioning of two +1 topological defects (due to Poincare’-Hopf index theorem for a sphere) at the top and the base of the bud. However this requirement of total +2 charge for a genus zero surface can be satisfied only when the bud detaches from the main body and not necessarily for a partial bud which is topologically connected to the basal surface. b) In ref[14] the bud formation (or narrowing of the neck) also requires a line tension at the neck where one type of lipid, constituting the bud, forms a raft like patch in the background of a second type of lipid. In our simulation the membrane is homogeneous or uniformly covered with one type of surface protein and therefore a case for line tension does not arise.

## Appendix

The divergence and curl on a curved surface can be found by using covariant derivatives. In this appendix, we outline the calculations for the derivation of Euler-Lagrange equations for cone and cylinder.

To write the Euler-lagrange equation, we need to calculate the expression for the frank free energy, which invols the calculation of divergence and curl on the 2D curved-surface. So, we define the following quantities which we’ll need for the derivation of curl and divergence [14]

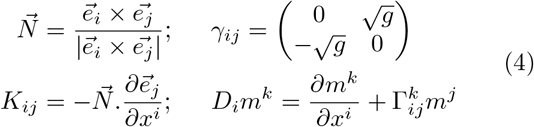

Where, 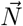 is the unit normal to the surface, *γ*_*ij*_ is the antisymmetric tensor, *g* is the determinant of the metric tensor matrix, and *K*_*ij*_ is the curvature matrix. *D*_*i*_ is the covariant derivative on the curved surface and 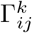 is the Christoffel symbol given by,

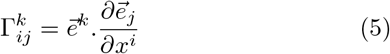

Now, the divergence and curl on the surface of a cone can be written as [14]

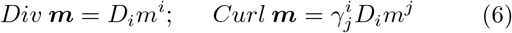

## Euler-Lagrange equation for Cone

The points on the surface of the cone can be represented by the three dimensional position vector 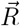. As we’ll be working on the surface of the cone, we can replace the cylindrical coordinate variables *z* and *r* as a function of *l*. From Fig.7, we can write *r*(*l*) = *l* sin(*α*), and *z* = *l* cos(*α*). So, the position vector can be written in cylindrical coordinate as

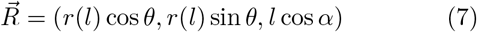

From the position vector we can construct a covariant basis from the expression 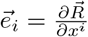, where *i* = 1, 2. Then the components of the metric tensor can be found from these basis vectors by using the definition 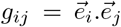. So, for conical surface, the metric tensor and curvature matrix(from eq.4) can be written as follows

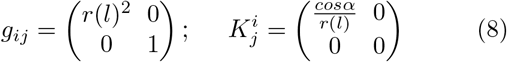

Now, using the curvature matrix we can write the helfrich-Prost term

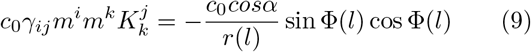

The components of 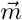 along 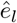 and 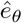 are *m*_*l*_ = cos Φ and *m*_*θ*_ = sin Φ, respectively. Now, using the expressions for divergence and curl in Eq. 6 we can write,

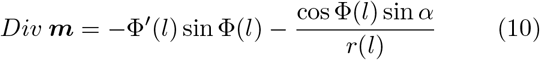

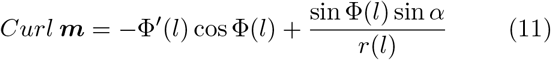

Prompted by the chiral nematic patterns found in our simulations (see Fig.3–6) we assume azimuthal symmetry, i.e., Φ is independent of *θ*. Using Eq.4,6 we can now write the frank free energy,

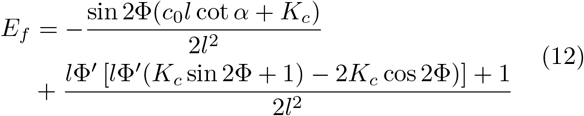

The corresponding Euler-Lagrange equation is

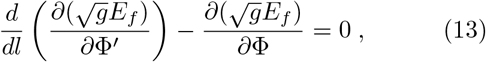

which yields,

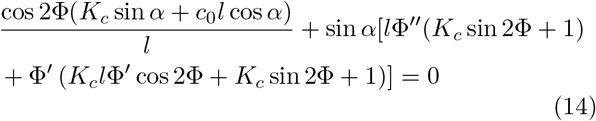

Numerical solutions of this equation are discussed in the main-text.

## Euler-Lagrange equation for Cylinder

**Figure 11:**
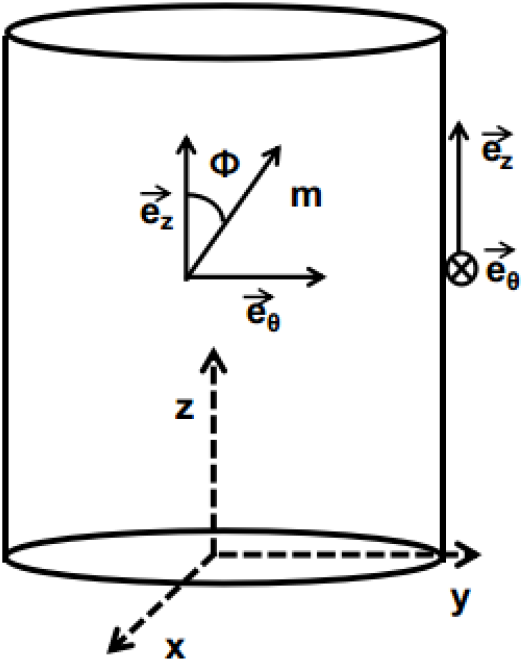
Schematic representation of cylinder and its basis vectors. The nematic director makes angle Φ(*z*) with the basis vector 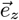, while 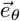 is the unit vector along the azimuthal direction.

Following the procedure described in the previous section, we proceed to compute the Frank free energy, starting with the position vector 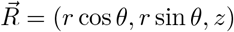,

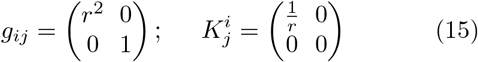

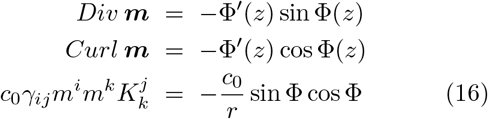

Assuming azimuthal symmetry, as before, Φ is a function of *z* only. Now using the above expressions, the Frank free energy reads

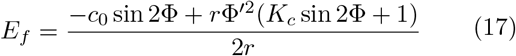

Using Eq.13, with Φ as a function of *z*, we can write the EL equation for a cylinder

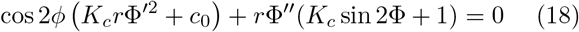

Note that, each of the above expressions for cylinder can be derived from the respective expressions for the cone, in the limit *α* → 0, *l* → ∞, keeping *r*(*l*) = *l* sin *α* finite.

## Acknowledgements

AA and AS acknowledge Science and Engineering Research Board(SERB), India(Project No. CRG/2019/005944) and IRCC-IIT Bombay for financial support. AB acknowledges the financial support by UGC, India and IRCC-IIT Bombay, and GK thanks NPDF-DST, India for financial support.

